# Even well-practiced movements benefit from repetition

**DOI:** 10.1101/2021.12.21.473484

**Authors:** Katrin Sutter, Leonie Oostwoud Wijdenes, Robert J. van Beers, W. Pieter Medendorp

## Abstract

Professional golf players spend years practicing, but will still perform one or two practice swings without a ball before executing the actual swing. Why do they do this? In this study we tested the hypothesis that repeating a well-practiced movement leads to a reduction of movement variability. To operationalize this hypothesis, participants were tested in a center- out reaching task with four different targets, on four different days. To probe the effect of repetition they performed random sequences from one to six movements to the same target. Our findings show that, with repetition, movements are not only initiated earlier but their variability is reduced across the entire movement trajectory. Furthermore, this effect is present within and across the four sessions. Together, our results suggest that movement repetition changes the tradeoff between movement initiation and movement precision.

**New & Noteworthy:** Professional athletes practice movements that they have performed thousands of times in training just before it is their turn in a game. Why do they do this? Our results indicate that both initial and endpoint variability reduce with repetition in a short sequence of reaching movements. This means that even well-practiced movements benefit from practice.

## Introduction

Variability is an inevitable part of all our movements. When learning a new movement, variability is initially high, but practice makes our movements more precise, i.e. less variable (1, 2). Practice is not only useful for learning a new movement, it seems also beneficial for already mastered movements. For instance, professional golf players practice their swing without hitting a ball before making the actual swing. Why are experts still practicing?

The benefits of movement repetition have been mostly described in terms of a reduction in movement reaction time (RT) (3, 4). However, for movements requiring high precision such as hitting a golf ball it is unlikely that practice before the real hit is motivated by a reduction of RT. A more plausible explanation may be that the benefit of repeating such a movement is that it makes the subsequent movement itself more precise. Yet, it remains unclear whether an expert, who has practiced the movement to perfection during training, would perform a more precise movement due to a single or a few practice swings just before it.

To answer this question it is important to acknowledge that movement precision is limited by noise in the sensorimotor system. Noise arises at all stages of sensorimotor processing, not only at the central planning stages but also peripherally, during movement execution (5–8). Variability related to movement planning (“planning noise”) arises from stochastic fluctuations in central neural activity (6, 9, 10); variability related to movement execution (“execution noise”) arises in the motor periphery and may be determined by factors such as the basic physiological organization of motor-unit pools and their muscle fibers (2, 6, 7, 11, 12).

From studies on trial-by-trial motor learning, it is known that errors in previous movements can be used to modify planning of future movements (6, 13). Computational studies have emphasized the different roles of planning and execution noise in correcting for previous movement errors when planning a new movement (6, 14, 15). In support, Cheng and Sabes (16) demonstrated empirically that both noise sources contribute comparably to the variability in the trial-by-trial dynamics of reach adaptation to shifted visual feedback. Van Beers (6) modelled and measured the pattern of endpoint error reduction for repeated movements to the same target based on the involvement of both noise sources. In their model, movement corrections from one trial to the next are made by correcting relative to the planned aim point (the point where the planned movement would end if it were generated in the absence of execution noise) of the previous movement. Although this model was originally not intended to describe how movement variability evolves over repeated movements, here we will test its additional prediction that movement variability reduces after each repetition (see Methods for further details).

To test how movement history affects current movement variability, we used a center- out reaching task in which participants had to move to the same target 2 to 6 times in a row. Since movement variability depends on RT (shorter RT leads to higher initial variability, see 17) and since RT reduces when movements are repeated (3, 4), the effects of repetition and RT on initial, evolving and final movement variability are not easily isolated. To allow us to disentangle their effects, we created a wide range of RTs for each repetition. That is, in half of the trials, we accompanied the onset of the visual stimulus with an auditory beep, known to reduce RT (18).

Since our experiment consisted of multiple sessions on different days, our approach allowed us to also measure the effects of repetition on movement variability on the longer time scale of days. Studies on motor skill acquisition and motor habits predict that across sessions learning performance is enhanced and reaches a plateau (19, 20). Our experiment allowed us to test two scenarios. If immediate history of movement does not enhance performance after elongated practice, this would lead to an extinction of the repetition effect in later sessions, which would suggest that professional golf players’ precision does not improve after a single practice swing. However, if practice on both long (session) and short (trials) timescales determines the precision of movement, a repetition effect should be present also in later sessions, suggesting that professional golf players do benefit from a single practice swing.

Our results show that even for well-practiced movements reach endpoint variability reduces with repetition, congruent with the model’s predictions. Moreover, a repetition effect persists across sessions indicating that even well-practiced movements benefit from repetition. Furthermore, we show that repetition reduces both variability and RT at the same time, thereby changing the tradeoff between movement initiation and movement precision.

## Methods

Details about the setup and methods used to measure and analyze reach kinematics as well as the general paradigm have been described extensively in Sutter et al. (17). Here we provide only a brief summary; part of the data (data from non-repeated movements) has been described in Sutter et al. (17) as well.

## Participants

We recorded 33,000 reaching movements of 11 participants (8 women, age 22-29 years), who all signed a written informed consent form and were naïve to the aim of the study. All participants self-reported to be right-handed, to have no motor deficits or neurological condition and had normal or corrected-to-normal vision. Participants received €50 for participation. This study was approved by the ethics committee of the Social Sciences faculty of the Radboud University in Nijmegen, The Netherlands.

### Experimental setup

Participants performed reaching movements with their right arm in the horizontal plane holding a planar robotic manipulandum (vBOT; 21). The forearm of the participant rested on an air sled that allowed frictionless movements. There was no direct visual feedback of the arm due to a semi-silvered mirror that covered the arm. Visual stimuli were presented on an LCD monitor (model VG278H, Asus) that was suspended above and viewed via a mirror. The refresh rate of the monitor was 120 Hz. A photodiode was used to measure the timing of the visual stimuli. The position of the robot’s handle and the photodiode output were sampled at 1000 Hz. Auditory stimuli were presented via over-ear headphones. The auditory stimuli were white noise beeps of 90 dB with a duration of 100 ms.

### Experimental Paradigm

Participants were instructed to make a single fast and accurate reaching movement from the home position (white disc, radius 0.4 cm, ∼30 cm in front of the participant’s midline) to the target (gray disc, 0.4 cm radius). The target was presented at one of four locations, 10 cm outwards from the home position at 45□, 135□, 225□ or 315□ from the forward direction. The trial started once the participant moved the cursor to the home disc aided by visual feedback about the handle (i.e., hand) position (red disc, 0.35 cm radius) (Figure 1A). After a random delay of 500 – 1000 ms, a visual target appeared and the participant had to reach out to the target. There was no visual feedback about the hand position after the cursor left the home disc until the online estimate of its speed dropped below 0.5 cm/s. Participants received a score between 0 and 100 points after each trial. The score was determined by the distance from the center of the target at the position where the speed dropped below 0.5 cm/s. A score of 0 points was given if the online estimate of the movement time or the reaction time was longer than 1000 ms.

**Figure 1.**
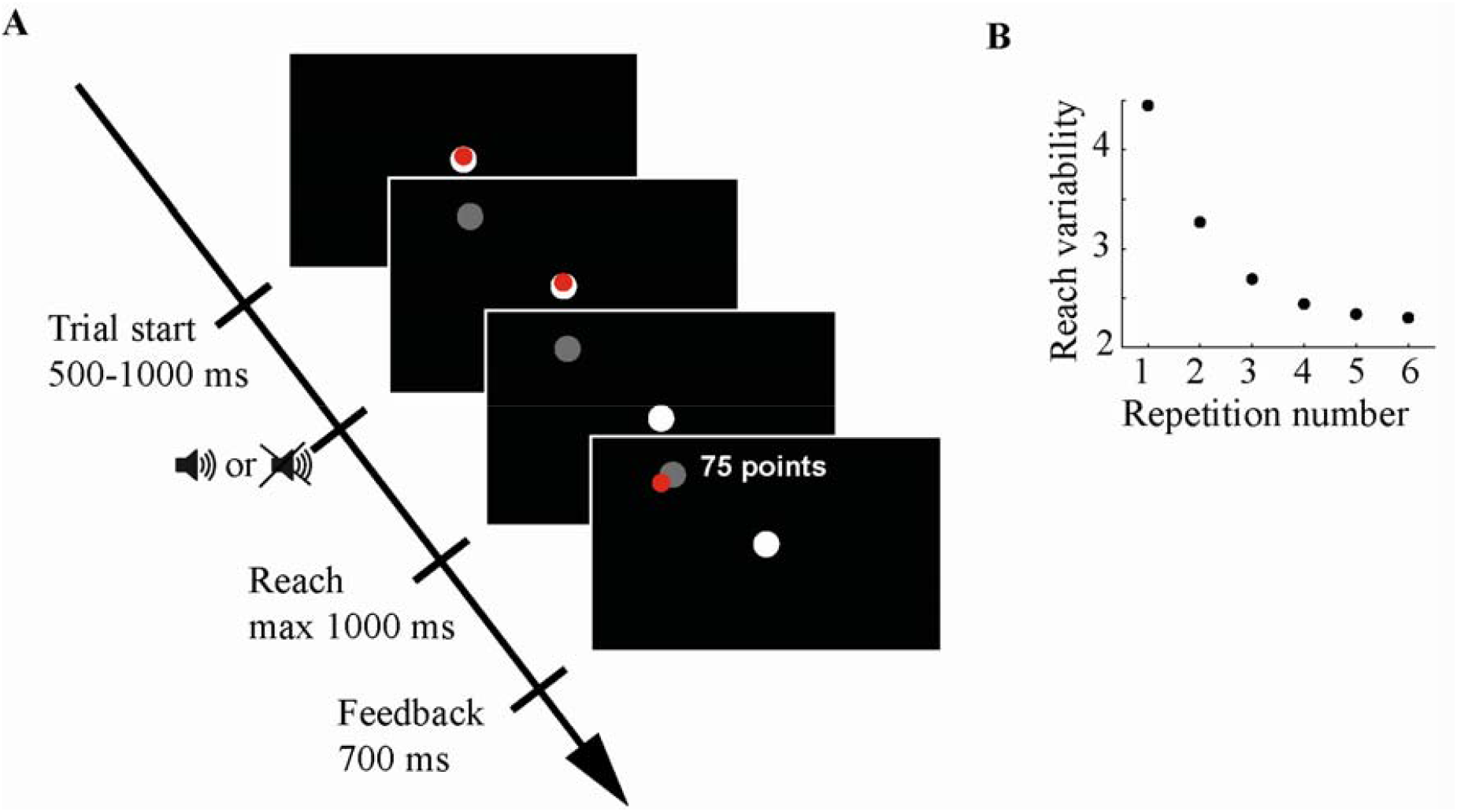
**A**. Timeline of the experiment. A trial started when the hand cursor was in the home disc; the target appeared 500-1000 ms later. In 50% of the trials, an auditory beep was played at target onset. The cursor disappeared upon leaving the home disc and appeared again when the velocity dropped below 0.5 cm/s. Participants received a score based on the distance between the cursor and the target. **B**. PAPC model predictions about reach endpoint variability with repetition.

To study variability in subsequent repetitions to the same target, a target could be repeated up to 6 times. Although the first trial in a sequence is formally not a repetition, we will refer to such trials as repetition 1, and similar for later trials. There were 80, 40, 20, 10 and 10 sequences of respectively 2, 3, 4, 5 and 6 repetitions to each target, yielding a total of 1880 trials. The different sequences were ordered randomly such that the same target could not be used in successive sequences. 280 single trials to each target were randomly inserted in between the sequences. Consequently, the proportion of trials that had the same target as the previous trial was about 50%. Results of the first trial in the sequence are reported in Sutter et al. (17).

To be able to distinguish between the effects of repetition per se and those of RT on variability in subsequent repetitions, in randomly selected trials (50% of total) the target onset was accompanied by an auditory stimulus. These trials are referred to as beep trials; the trials without auditory stimulus are referred to as no beep trials. Trials in the same sequence of repetitions to one target had the same type of auditory stimulus (beep or no beep).

Participants performed a total of 3000 trials, tested in 4 separate sessions of 12 blocks of about 62 trials. There was a break of at least 20 s between the blocks. Sessions took place on different days with a maximum of 5 days between sessions. All sessions started with a practice block in which the targets were organized in cardinal directions in order to avoid response facilitation (3). There were 40 practice trials in the first session and 20 in other sessions. The duration of the whole experiment was 270 min.

### Data analysis

Positional data analyses were performed with Matlab 2018b (MathWorks). Statistical data analyses were performed using SPSS Statistics 25. Position data of the robot handle were filtered with a 5^th^ order low-pass Butterworth filter with a cutoff frequency of 20 Hz. Reach onset was defined as the first time point when the hand cursor left the home disc. Reaction time was determined as the time from the onset of the visual target to reach onset. Reach direction was defined as the angle of the cursor (i.e., hand) relative to a straight line through the target, both referenced to the point where the hand left the home position.

Reach offset was determined using the multiple sources of information method (22) as the most likely time point considering objective functions of four different components: 1) time, shorter time since movement onset was given a higher value; 2) distance, longer distance from home was given a higher value; 3) velocity, lower velocities received a higher value; and 4) acceleration, positive acceleration was given 0 value. Movement time was defined as the time between reach onset and offset. Position data were resampled between reach onset and reach offset to contain 100 samples per trial. Reach variability for each timepoint was determined by first calculating the standard deviation of the reach direction across trials to the same target, separately for each trial type (beep or no beep) and repetition, and then taking the mean of the variability across the four targets. Initial reach variability was defined as the reach variability at the first time point after the hand cursor left the home disc (reach onset). Endpoint variability was defined as reach variability at reach offset.

Trials were removed from further analysis if the distance between the target and the endpoint of the movement was larger than 5 cm (n = 1), if the reaction time was above 1000 ms or below 100 ms (n = 16), or if reach direction deviated at any point of the movement more than 45 degrees from the initial reach direction (n = 164).

To validate pooling over beep and no beep trials (see below), we tested if the beep affected initial and endpoint variability. We calculated the mean initial and endpoint variability of trials within a 30 ms time window around the subject’s mean RT in that repetition. Thereafter we performed a 3-way ANOVA with movement phase (initial and endpoint), condition (beep and no beep) and repetition (1, 2, 3, 4 and 5) as within subject factors. In this analysis we included trials from sequences up to 5 repetitions, because not all participants had trials for the 6^th^ repetition in this particular time window.

We tested whether there were any session effects on the endpoint variability with a repeated measures ANOVA with session (1, 2, 3 and 4) and repetition (1, 2, 3, 4, 5 and 6) as within subject factors. We report partial eta squared (η_p_^2^) as a measure of effect size for main effects and interactions of the repeated measures ANOVAs and Cohen’s d for paired t-tests. Where appropriate, we applied Greenhouse-Geisser corrections.

### Model

We used the planned aim point correction (PAPC) model proposed by Van Beers (6) to quantify how trial-by-trial planning corrections affect the movement endpoint variability in a sequence of repeated movements. This model assumes that motor plans are generated centrally. If a motor plan in trial *t* would drive the movement in the absence of noise in movement execution, the movement would end at location 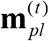, the *planned aim point*. However, since there is noise in movement execution, its effect 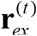 needs to be added to the planned aim point in order to obtain the endpoint **x**^(*t*)^ of movement *t*:

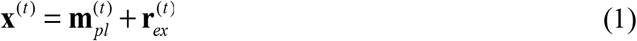

The error **e**^(*t)*^ in movement *t* is the difference between the endpoint and the target location **x**_*T*_:

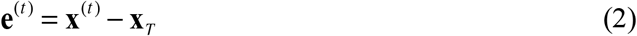

This error is used to improve planning of the next movement **m**_*pl*_ ^(t+1)^ by correcting the planned aim point of the previous trial by a proportion *B*, the *learning rate*, of this error, while adding the effect of noise in movement planning,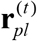 :

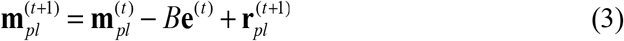

The random effects of planning and execution noise are drawn from zero-mean Gaussians with covariance matrices Σ _*pl*_ and Σ_*ex*_, respectively. Since the planning of the first movement to a target in a sequence is generally less precise (6), an additional random vector **r**_0_, drawn from a zero-mean Gaussian with covariance matrix Σ_0_, is added to the planned aim point of the first of a sequence of movements to a target. The various covariance matrices are parameterized as: Σ _*pl*_ = *w*Σ_*mot*_, Σ_*ex*_ = (1− *w*)Σ_*mot*_ and Σ_0_ = *k*Σ_*mot*_, where *w* is the proportion of the total amount of motor covariance that arises during planning, Σ_*mot*_ is a generic covariance matrix, and *k* is a scaling factor (see (6) for details).

We here derived an equation for how the covariance of movement endpoints depends on the repetition number *t* in a sequence of movements to the same target (see Appendix A for the derivation):

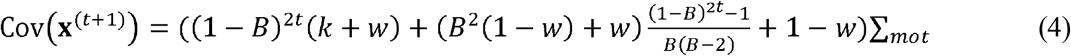

With the parameter values from (6) (*w* = 0.21, *B* = 0.39, *k* = 4, Σ_*mot*_ = 4 (in this study, we consider only a single dimension)), the PAPC model predicts that the reach endpoint variability reduces with repetition (Fig. 1B). The reason it predicts this reduction is that the variability is relatively high in the first movement of a sequence (as quantified by scaling factor *k*), and that the planning corrections based on observed errors lead to more accurate and therefore less variable planning of later movements.

## Results

In this study we tested how repeatedly reaching to the same target affects movement variability. In addition, we investigated how the effects of reaction time and repetition interact. To evoke a wide range of RTs necessary for this aim we used an accessory auditory stimulus in 50% of the trials. The RT of trials with a beep was on average 26 ms shorter (286 ± 29 ms, mean ± SD) than the RT of trials without a beep (313 ± 28 ms). To verify that the auditory stimulus did not affect the variability of the reach, we compared the variability of trials with and without a beep within a 30 ms time window around the participant’s mean RT in that repetition. There was no main effect of beep (*F*(1, 10) = 0.17, *p* = 0.90, η_p_^2^= 0.002) or any interaction effect with beep. This confirms that the accessory auditory stimulus did not affect the variability of the reaching movement, allowing us to pool the trials with and without a beep in our subsequent analyses.

Figure 2 shows how reach variability evolves when the movement unfolds during the first (red) and subsequent target repetitions. Both initial and reach endpoint variability reduce with repetition number. The reduction in variability with repetition persists from the start until the end of the reach. It can further be observed that repetition leads to a reduction of RT, as the curve shifts towards the left with repeats. Thus, complementary to our previous study (17), in which we found that in non-repeated movements a shorter RT causes higher initial variability, we observe that movement repetition results in shorter RTs but not at the expense of increased variability. We will now examine each of these findings in more detail, starting with the endpoint variability.

**Figure 2.**
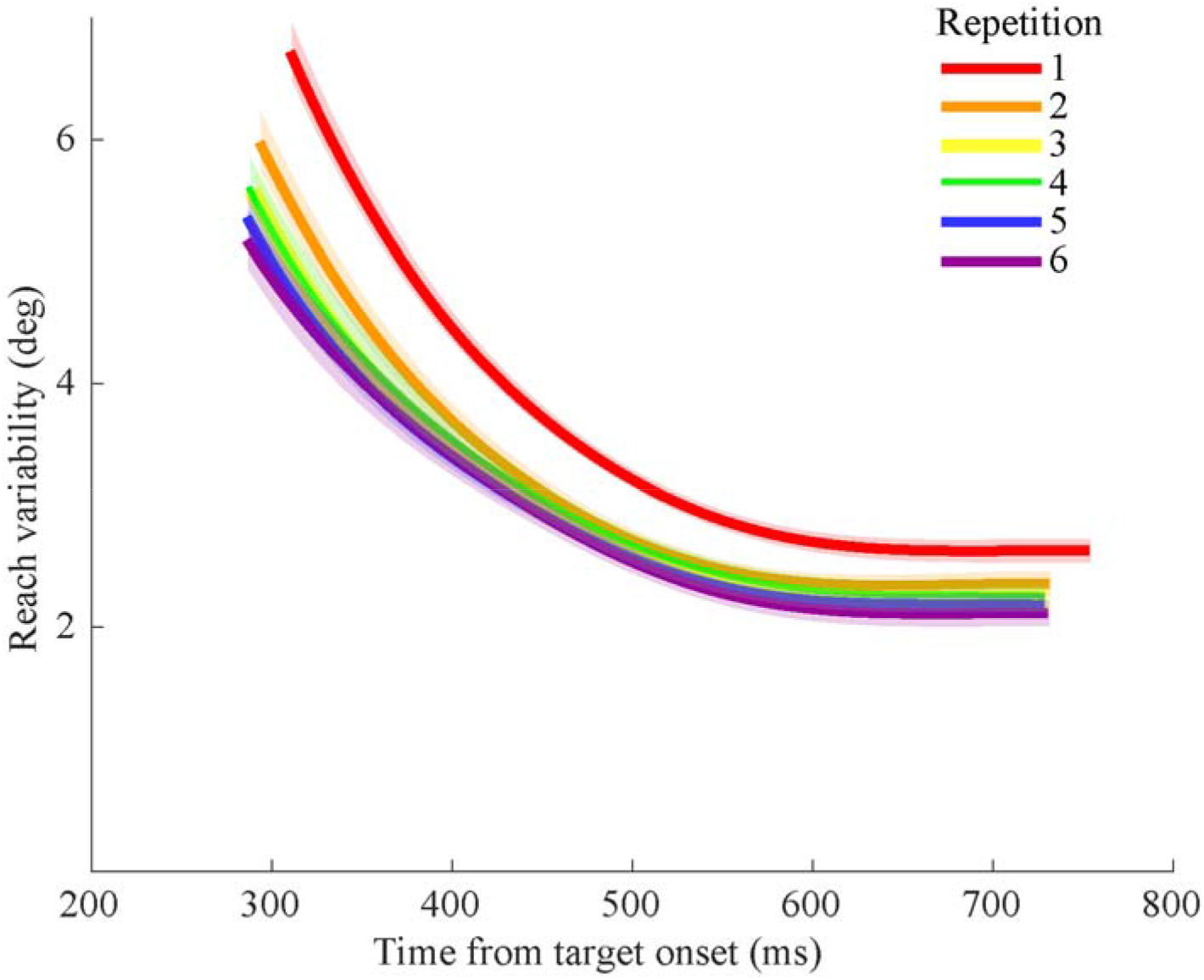
Reach variability as a function of time from target onset for different repetitions. The starting time of each line is the mean group RT of the respective repetition. The end time of each line is based on the average movement time (MT) of the respective repetition. Shaded areas represent SEM. Repetition 1 (in red) is formally not a repetition – it is the first trial of the sequence.

### Endpoint variability

To test whether endpoint variability reduces with repetition we performed a repeated measures ANOVA with repetition number (1, 2, 3, 4, 5 and 6) as the within subject factor. There was a significant effect of repetition on the endpoint reach variability (F(5, 50) = 15.43, p < 0.001, η_p_^2^ = 0.61). *Posthoc* comparisons revealed that endpoint variability was significantly reduced in the first 3 repetitions (all p < 0.036). Furthermore, to test the predictions of the PAPC model we estimated three model parameters based on the group average of reach endpoint variability in different repetitions. We fixed the fraction of planning noise in the endpoint variability (w) at 0.21 as established previously (6).

The model had an R^2^ of 0.79 (Figure 3A). These results indicate that variability reduces with repetition as predicted by the model, suggesting that there is trial-to-trial learning. The best estimates for the free parameters were: learning rate B = 0.33; variance of effect of motor noise ∑_mot_ = 3.48 deg^2^; and scaling factor for the covariance of the first movement in a sequence k = 0.62.

**Figure 3.**
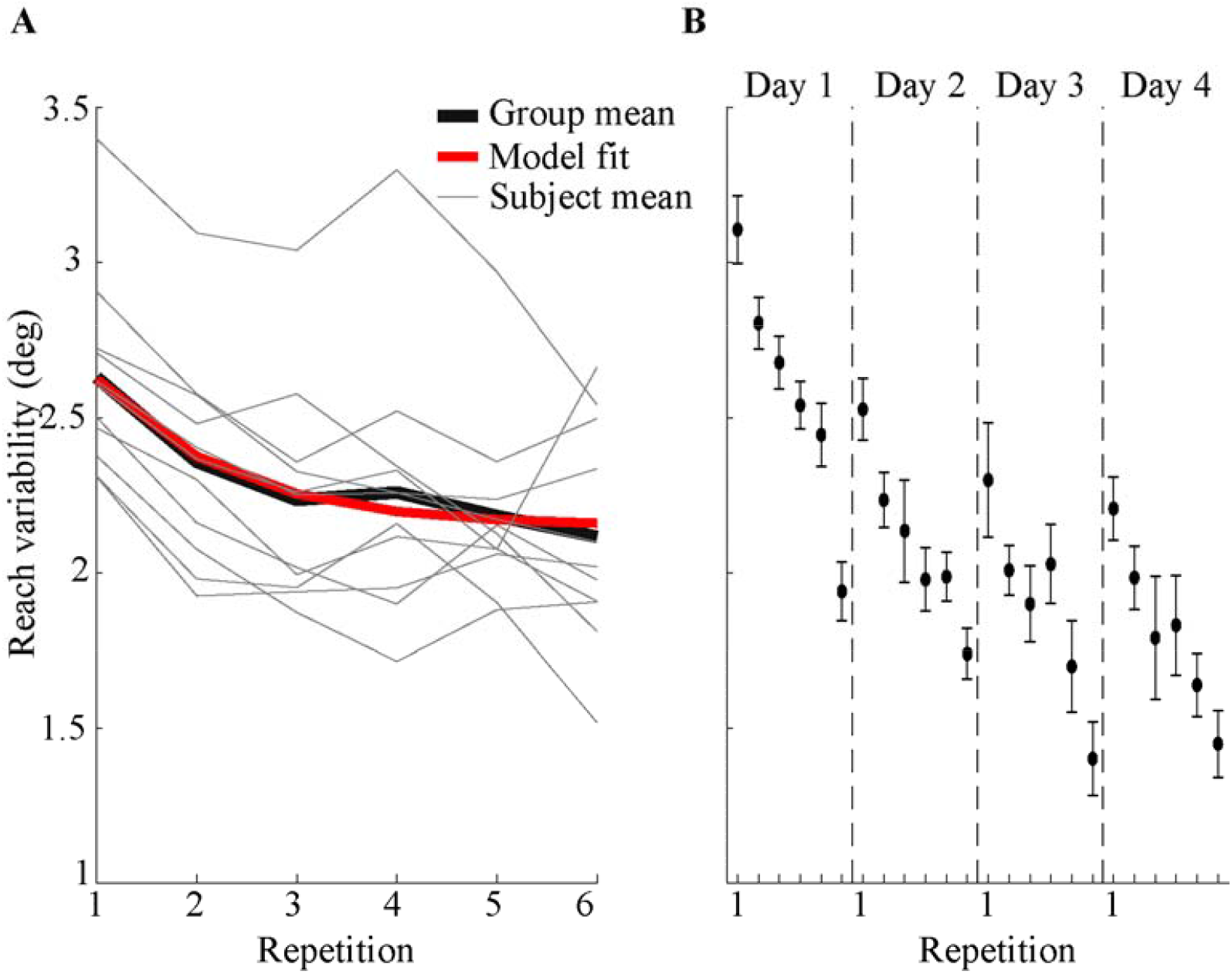
**A**. Reach endpoint variability and model fit. The thick black line represents the mean endpoint variability of the participants; the gray lines depict the individual participants. The red line represents the model fit. **B**. Reach endpoint variability on different days averaged across participants. Dashed lines indicate the separation between different days. Error bars represent the SEM.

### Initial variability

Figure 2 suggests that, as endpoint variability, also initial variability reduced with repetition. Performing a similar analysis on the initial variability as on the endpoint variability would however not be legitimate because, unlike the endpoint variability, the initial variability depends on the RT (17). We therefore first analyzed how the initial variability varied with both repetition number and RT. For this purpose, we used a median split per participant resulting in two bins with short and long RT trials. We then computed the initial variability for the short and long RT bins separately, and plotted the mean variability in the short and the long RT bins as a function of repetition number. As shown in Figure 4, initial variability depends on both on RT and repetition number. To formally test this, we performed a 2 × 6 repeated measures ANOVA with RT bin (short, long) and repetition number (1, 2, 3, 4, 5 and 6) as within subject factors. Confirming the observations, we found a significant effect of the RT bin (F(1,10) = 10.27, *p* = 0.009, η_p_ ^2^ = 0.51). Short RT trials were on the average more variable than long RT trials (6.09 ± 0.23 and 5.79 ± 0.21 deg). We also found a significant effect of the repetition number (F(5,50) = 47.50, *p* < 0.001, η_p_^2^ = 0.83) and a significant interaction effect (F(5,50) = 17.11, p< 0.001, η_p_ ^2^ = 0.63), which was driven by the higher repetition effect within the short RT trials when compared to the long RT trials.

**Figure 4.**
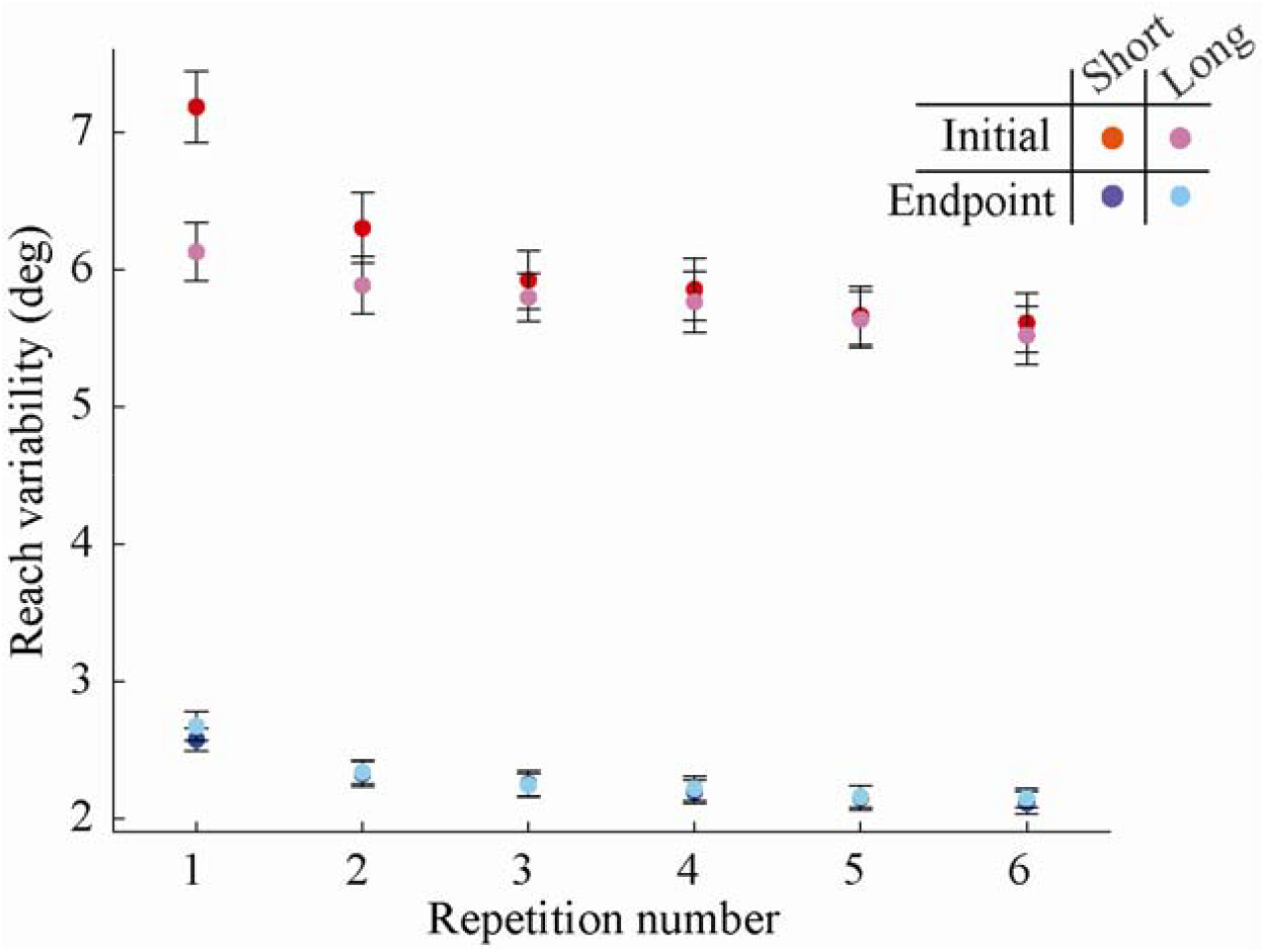
Initial and endpoint reach variability of short and long RT trials as a function of repetition number. Error bars show the SEM.

To test whether the RT also affects reach endpoint we performed the same statistical analysis on endpoint variability and plotted the endpoint variability in short and long RT bins on Figure 4. We did not find a significant effect of RT (F(1,10) = 2.0, p = 0.19, η_p_^2^ = 0.17).

This suggests that only the initial variability is affected by an interaction between repetition number and RT.

To determine the effect of repetition number on initial variability while circumventing the complex interaction between reaction time and repetition number, we compared trials from different repetitions in a 30 ms time window around each participant’s mean RT. Figure 5 shows how reach variability reduced with repetition in the first five movements to the target, even if the trials are from the same RT window (initial variability in increasing order of repetition: 6.50 ± 0.78, 5.75 ± 0.85, 5.40 ± 0.44, 5.42 ± 1.08 and 4.95 ± 0.78 deg). We did not include the trials of the 6^th^ repetition, because not all participants had trials within the time window (see Methods). A repeated measures ANOVA revealed a significant main effect of repetition (F(4,40) = 7.32, *p* < 0.001, η_p_ ^2^ = 0.42) on initial variability. Figure 5 illustrates that this effect was preserved until the end of the movement. This analysis confirms that movement repetition reduces initial variability, even for the trials of different repetitions within the same RT range.

**Figure 5.**
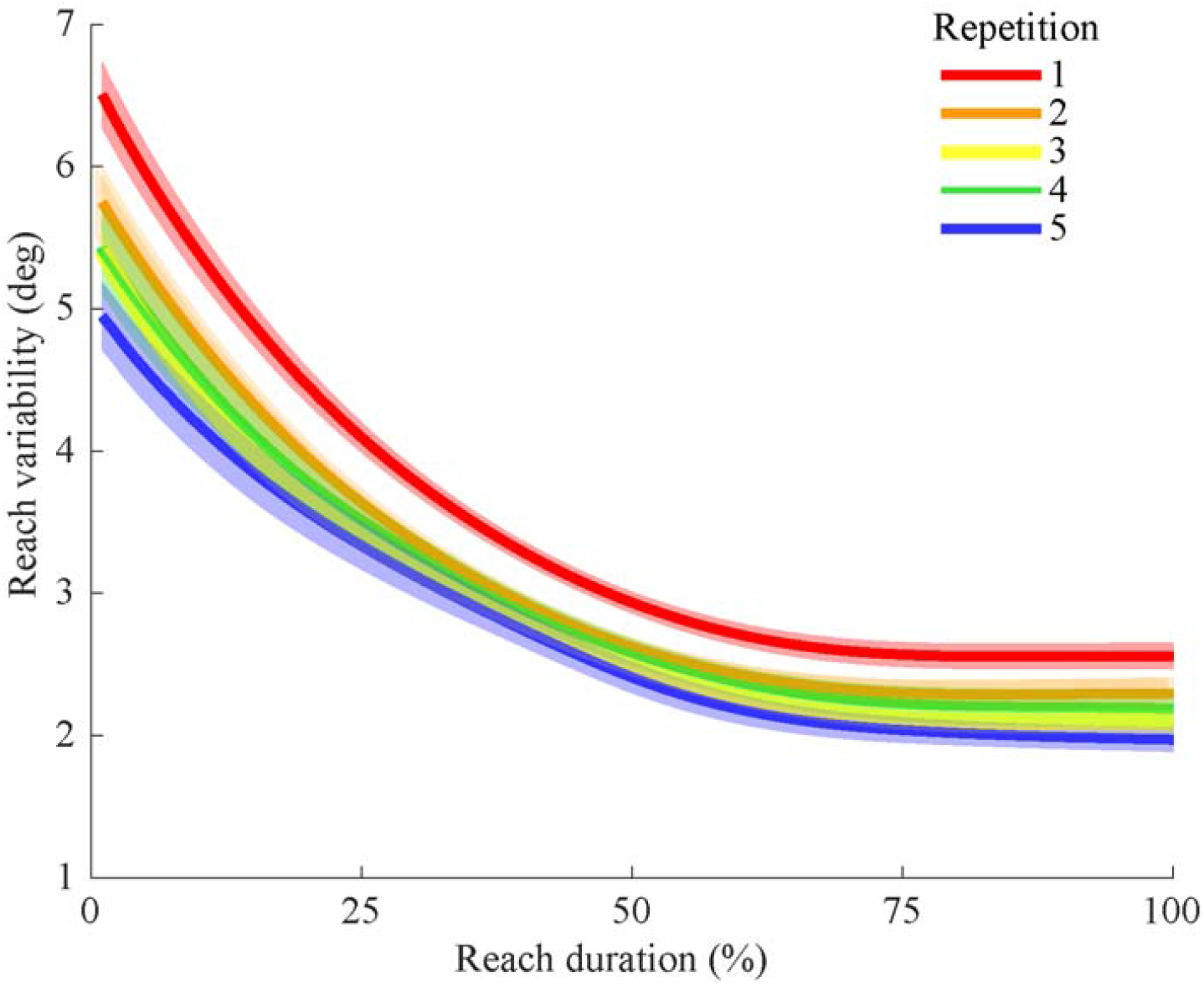
Reach variability as a function of reach duration for trials within a 30 ms time window around the participant’s mean RT for repetitions 1-5. Shaded areas represent SEM.

### Session effects

We next examined if there were any session effects on endpoint variability and whether the repetition effect persisted over sessions (Figure 3B). Average reach endpoint variability reduced over sessions (variability was 2.59 ± 0.13, 2.10 ± 0.07, 1.89 ± 0.07 and 1.82 ± 0.09 deg (mean ± SEM) in sessions 1 to 4). A 4 × 6 repeated-measures ANOVA on reach endpoint variability with session (1, 2, 3 and 4) and repetition number (1, 2, 3, 4, 5 and 6) as within subject factors confirmed that the session effect was significant (F(3,30) = 40.08, p < 0.001, η_p_^2^ = 0.80). In line with our main hypothesis, we also found a significant effect of repetition number (F(5,50) = 71,40, *p* < 0.001, η_p_^2^ = 0.88). There was no interaction between session and repetition number (F(15,150) = 1.21, p = 0.27, η_p_^2^ = 0.11). This confirms that repeatedly moving to the same target results in lower endpoint variability both within and across sessions. Importantly, repetition reduced variability in each session.

### Reaction time

A previous study (3) has shown that repetition reduces RT. We replicate this finding: Figure 6 illustrates that the group mean RT reduces with repetition. A one-way repeated measures ANOVA with repetition number as the within subject factor revealed a significant effect of repetition number on RT (F(5,50) = 45.6, *p* < 0.001, η_p_ ^2^ = 0.82). *Posthoc* tests revealed that RT significantly reduced in the first 3 repetitions (p < 0.0004).

**Figure 6.**
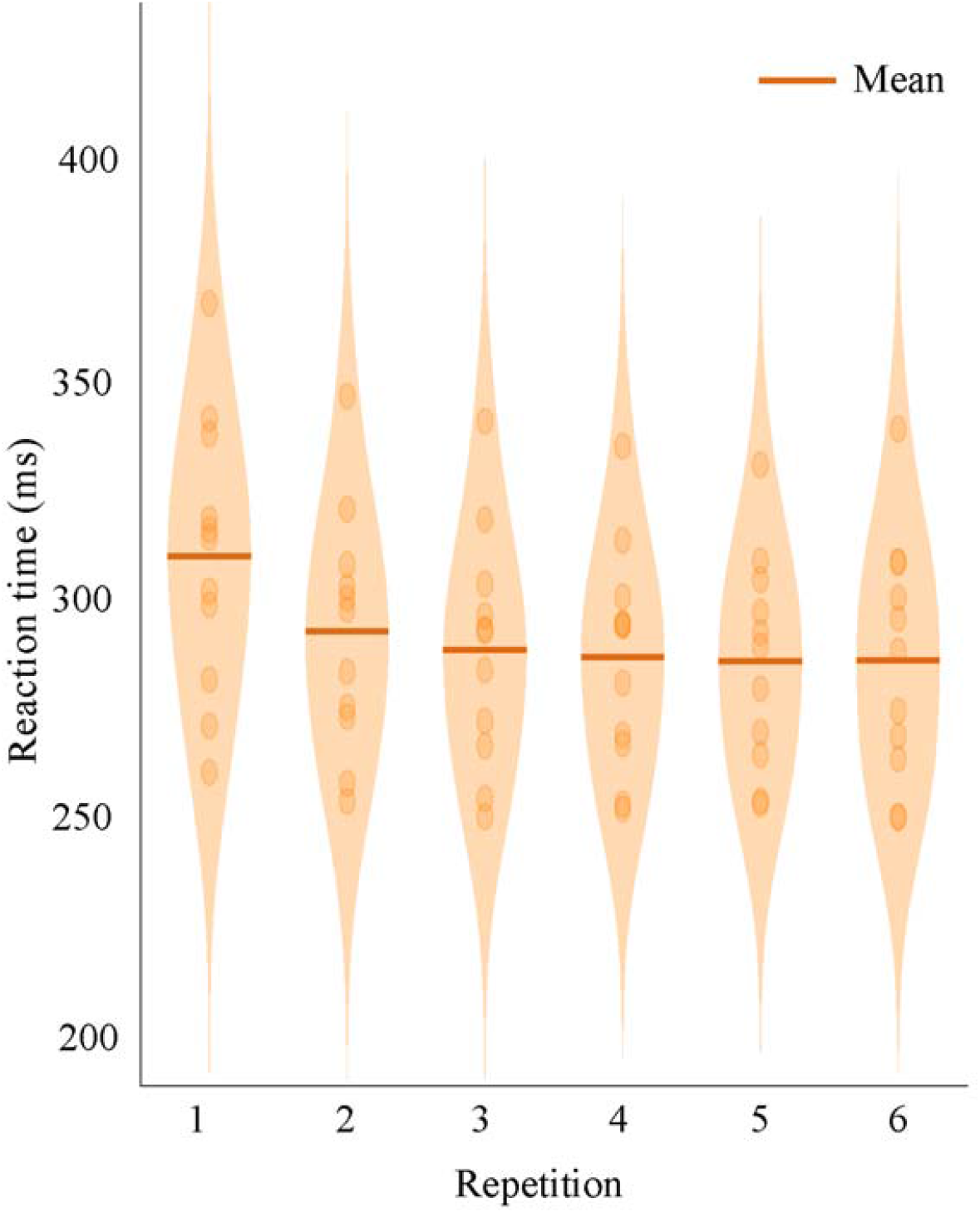
Violin plot of mean reaction time as a function of movement repetition. Values for individual participants (dots) and the group mean (bold line) are also indicated.

## Discussion

We investigated whether repeating well-practiced movements affects movement variability. In a center-out reaching task, participants made movements towards 4 targets that could be repeated up to 6 times in a sequence. We show that repeating even well-practiced movements reduces the variability with which the movement unfolds (Fig 3A, Fig 5). We found that variability reduced not only within a sequence of repetitions, but also across experimental sessions (Fig 3B). Furthermore, we show that initial variability is affected by both repetition and RT, whereas the endpoint variability is only affected by repetition (Fig 4, Fig 5).

Endpoint variability reduced in a sequence of movements with each repetition. These observations are in line with the prediction of the PAPC model by van Beers (6). The predictions of this model for precision improvement are comparable with the effect that is called warm-up decrement in the sports sciences. Warm-up decrement entails that people have to retune or recalibrate their motor systems for reaching the peak performance at a task (23, 24).While the warm-up decrement has been known for a long time (24), there is little research and understanding of its origin. A recent examination by Wunderlich and colleagues (25) of throwing data of professional darts players showed similar improvements in accuracy as we show with reach variability. In professional darts, players take turns throwing three darts.

Wunderlich and co-workers show that throwing accuracy is the lowest in the first throw of each turn. Similar results have also been reported for free throws in basketball (26). Data from the NBA seasons between 2006 and 2016 show that with double free throws the success rate was higher for the second than for the first throw, and with triple free throws the success rate increased with each successive throw. Wunderlich and colleagues (25) suggest that as the warm-up decrement occurs within the short time scale of seconds it is unlikely that the changes in performance are due to changes in arousal, attention or other bodily or mental states. They suggest instead that the observed increase in accuracy is due to fine-tuning of visuomotor calibration. They hypothesize that moving away from the dartboard in between turns leads to loss in visuomotor calibration with respect to targets defined in an external frame of reference. We propose that the visuomotor calibration that these authors (25) describe involves trial-by-trial learning. Predictions of the PAPC model and of similar models describing the dynamics of trial-by-trial learning (14, 15) are in line with the studies on warm-up decrement and bring a modeling perspective in understanding how the brain uses previous movements to improve planning of future movements.

While repeating a movement can lead to performance improvements, it can also raise a bias for the upcoming movements. Verstynen and Sabes (27) showed that repeating one target location more often than the others, leads to reduction in movement variability at the cost of an increase in movement direction bias. In particular, the precision of the reaches to the repeated target became more precise, but the reaches to the other targets became biased towards the repeated target. Similarly, in a reaching task with many possible solutions, passive movements to the target along a certain trajectory bias the upcoming movements towards that trajectory (28). This phenomenon is known as use-dependent learning (27–29). An important difference between those studies and ours is that we did not vary how often had to be moved in different directions, as all targets were presented equally often. Furthermore, we show a variability reduction on the timescale of one or a few trials as opposed to timescales of tens of trials in the studies on use-dependent learning. It is therefore unlikely that the short-term effects we found reflect use-dependent learning.

We have previously shown that movement preparation time determines initial movement variability: the shorter the RT, the higher the initial variability (17). We established this tradeoff based on the first trial of the sequence, hence isolated the RT effects from repetition effect. The current study indicates that repeating a movement to the same target changes this tradeoff: movement repetition leads to both RT reduction and variability reduction.

We further show that repetition reduces variability during the entire movement, whereas preparation time has an effect on movement variability only for the initial phase of the movement (17). The trial-by-trial corrections driven by previous errors lead to improvements for the entire movement, because the entire movement plan can be adjusted. The effect of preparation time that we studied previously (17) is not governed by error-driven corrections, so an increased preparation time does not carry a signal to learn from that could affect the movement endpoint. Importantly, also in the current study we do not find an effect of preparation time on the endpoint variability of movements in different repetitions.

We show that movement variability also reduces across days, which may be interpreted within theories about multiple timescales of motor learning (30). Experimental motor learning tasks that take minutes, an hour or a few days to learn are often modeled using a fast and a slow process. However, many tasks (e.g. golf) require long practice before they are mastered and frequent practice is needed for maintaining the attained level of skill. In contrast, some skills, like riding a bicycle, seem to be impossible to forget. Recent behavioral and computational studies show that there are (at least) three learning processes with different timescales: fast, slow and ultraslow (31, 32). These processes are in charge of the balance between retention and forgetting. While the PAPC model only has one timescale (defined by the learning rate), we assume that the scaling factor *k* for the covariance of the first movement in a sequence reduces across days, as participants got more experienced in making these movements over days. Therefore, although our experimental task is relatively easy and spans only 4 different experimental days, we expected to see a decline in variability across the days. This explains also why the scaling factor for the covariance of the first movement in a sequence in the current experiment was considerably lower (scaling factor: 0.62) than in the van Beers (6) experiment (scaling factor: 4). In the current experiment, participants experienced many “first trials” in a sequence, while in the van Beers study there was only a single sequence of 30 sequential movements to each target.

Does the repetition effect require actual movement execution or is movement pre-planning also sufficient? The difference between only planning or executing a movement has been studied in the context of motor adaptation. It is known that learning two opposing force fields at the same time is a challenge. Sheahan and colleagues (33) show that planning without executing different follow-through movements is sufficient for learning opposing perturbations simultaneously. Furthermore, Kim and colleagues (34) have recently shown that motor learning in the context of motor adaptation occurs also if the previous movement is withheld. These observations are in line with studies showing that motor imagery improves performance in sports (35, 36). In addition, Ariani et al (37) showed that movement repetition can also affect reaction time (RT), movement time (MT) and movement accuracy. In a go/no-go paradigm with a discrete sequence production task, they showed that the repetition effect on RT and MT was present only when the movement was actually repeated (37). On the contrary, they also showed that accuracy was improved even when the previous trial was a no-go trial, such that the movement was only pre-planned, but not executed. However, as accuracy and variability represent different aspects of a data distribution, this does not imply that only pre-planning is sufficient for a reduction in variability. It remains to be determined whether movement planning is also sufficient for the repetition effect reported here.

Is there a difference in the repetition effect between repetition with error feedback (e.g. dart player’s first throw) and without error feedback (e.g. the golfer practicing the swing without hitting the ball)? While in the latter case the golf player would indeed miss the information about where the ball would land, the sensory feedback about the performance of the swing is available and can be used to predict the landing location of the ball. Therefore one could reason that the feedback in that case is not missing, but of lower quality. Interestingly, the quality of the error feedback in a reaching task affects the learning rate: the more reliable the feedback, the higher the learning rate (38–40). On this basis, it can be expected that the repetition effect will be stronger when more reliable error feedback is available.

What are the implications of the present findings at the neural level? Recent neurophysiological studies with delayed reaching tasks suggest that neural control of movement is achieved by time-evolving changes in the population of neurons in movement-related brain regions. Such a dynamical systems perspective explains that the neural population activity modulates itself in the different phase of movement generation, traveling a trajectory along the optimal states necessary for movement planning, initiation and execution, all modulated by sensory feedback (41). Ariani and colleagues (37) studied movement preparation of sequential finger movements. Based on their findings of shorter inter-press-intervals with repetitions they suggest that at a neural level repetition leads to reaching the initial state faster and more efficiently by bypassing intermediate suboptimal stages (42). Our results would be in line with this notion, although neuronal measurements are needed for direct confirmation.

## Grants

This work was supported by TopTalent grant from the Donders Institute to K.S. and a grant from the Netherlands Organization for Scientific Research (NWO-VICI: 453-11-001) to W. P. M.

## Appendix

Based on the PAPC model, an equation for the endpoint covariance as a function of repetition number *t* in a sequence of movements to the same target can be derived. The covariance of the planned aim point in repetition *t* + 1 is:

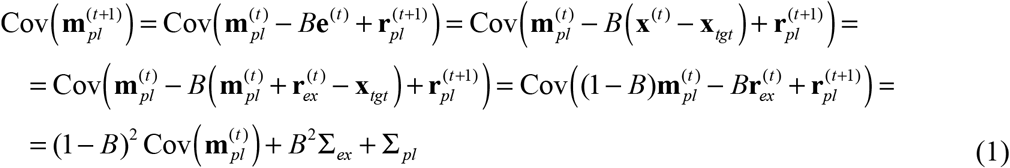

This is a recursive relation of the form 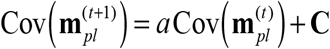. From this recursive relation, the following equality can be derived:

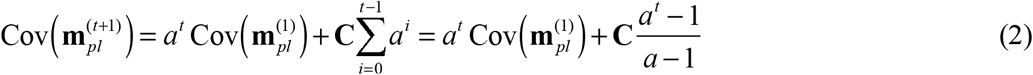

With 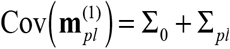 and *a* = (1− *B*)^2^, **C** = *B*^2^ Σ^*ex* +^Σ_*pl*_, it follows that:

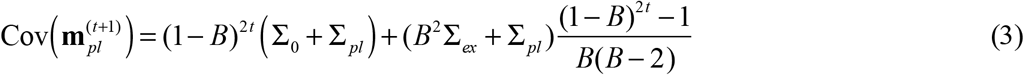

This is the equation for the covariance in the planned aim points. Since the endpoint differs only a random vector 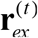 from this, the covariance of the endpoints is:

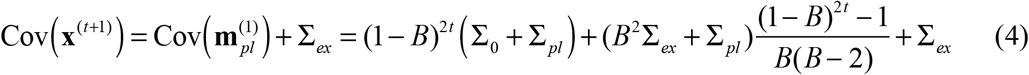

With the parameterization of the covariance matrices given above, this takes the form of equation 4 in the main text.

## Notes

### Competing Interest Statement

The authors have declared no competing interest.

